# Quantitative physiology and proteome adaptations of *Bifidobacterium breve* NRBB57 at near-zero growth rates

**DOI:** 10.1101/2022.09.06.506712

**Authors:** Angela Rocio Ortiz Camargo, Oscar van Mastrigt, Roger S. Bongers, Kaouther Ben- Amor, Jan Knol, Tjakko Abee, Eddy J. Smid

**Affiliations:** Food Microbiology, Wageningen University & Research, Wageningen, The Netherlands; Danone Nutricia Research, Utrecht, The Netherlands; Laboratory of Microbiology, Wageningen University & Research, Wageningen, The Netherlands

**Keywords:** retentostat, proteomics, chemostat, stringent response, metabolism, bifidobacteria

## Abstract

In natural environments, nutrients are usually scarce causing microorganisms to grow slow while staying metabolically active. These natural conditions can be simulated using retentostat cultivations. The present study describes the physiological and proteome adaptations of the probiotic *Bifidobacterium breve* NRBB57 from high (0.4 h^−1^) to near-zero growth rates. Lactose-limited retentostat cultivations were carried out for 21 days in which the bacterial growth rate progressively reduced to 0.00092 h^−1^, leading to a 3.4-fold reduction of the maintenance energy requirement. Lactose was mainly converted into acetate, formate and ethanol at high growth rates while in the retentostat lactate production increased. Interestingly, the consumption of several amino acids (serine, aspartic acid and glutamine/arginine) and glycerol increased over time in the retentostat. Morphological changes and viable but non-culturable cells were also observed in the retentostat. Proteomes were compared for all growth rates, revealing a down-regulation of ribosomal proteins at near-zero growth rates and an up-regulation of proteins involved in the catabolism of alternative energy sources. Finally, we observed induction of the stringent response and stress defence systems. Retentostat cultivations were proven useful to study the physiology of *B. breve*, mimicking the nutrient scarcity of its complex habitat, the human gut.

**IMPORTANCE:** In natural environments, nutrients are usually scarce causing microorganisms to grow slow while staying metabolically active. In this study we used retentostat cultivation to investigate how the probiotic *Bifidobacterium breve* adapts its physiology and proteome under severe nutrient limitation resulting in near-zero growth rates (<0.001 h^−1^). We showed that the nutrient limitation induced a multifaceted response including stress defence and stringent response, metabolic shifts and the activation of novel alternative energy producing pathways.

## Introduction

In natural environments, microorganisms are not exposed to optimal growth conditions (1). Often, nutrients are not readily available for consumption because of low concentrations and competing microorganisms. This causes a scarcity of nutrients and hinders microbial growth by inducing starvation (2, 3). Consequently, microorganisms enter a famine phase where growth becomes slow while they stay metabolically active. This forces bacteria to become energy efficient in order to survive (4, 5).

Several systems are used to study microbial physiology in the laboratory. The most common method is batch cultivation. Nevertheless, this model does not reflect well microbial life in natural environments (4, 6). In batch cultivations, all nutrients are initially abundant resulting in fast microbial growth. Nonetheless, in batch cultures, the concentrations of some nutrients may become too low to support further growth. To grow microorganisms under more controlled conditions, chemostat cultivation is used where the dilution rate determines the growth rate. Thus, fresh medium is continuously fed at a fixed rate into the bioreactor while there is a simultaneous removal of spent medium containing biomass, metabolic products, and non-used nutrients (7). However, growth rates in chemostat cultivations cannot reach those low levels usually observed in nature, which are close to zero (8).

A better system for the controlled cultivation of microorganisms at very low growth rates is the retentostat (9). The retentostat is a modification of the chemostat, in which the biomass is completely retained in the bioreactor using a filter in the effluent line (10, 11). In this way biomass accumulates reducing the available energy sources per individual cell resulting in a gradual decrease in growth rate that approaches zero (11). Therefore unlike the chemostat, the retentostat cultivation allows us to emulate the natural environmental conditions of microorganisms (8).

Retentostat studies have shown the importance of this system in elucidating the mechanisms involved in survival of microbial cells in environments with extremely low nutrient availability (4, 10, 12–17). In these conditions, microbial cells use different strategies to adapt to the nutritional stress in order to survive. Microbial cells make efficient use of the available energy by shutting down certain processes involved in growth, while activating others that would be advantageous for survival (12, 18). Hence, maintenance requirements are extremely reduced at near-zero growth rates (13, 16, 19). Concomitantly, shifts in the metabolism are seen where pathways are activated for the use of alternative energy sources (4). Changes in the morphology, the generation of viable-but-non-culturable (VBNC) cells and improved robustness have also been reported as survival strategies at near-zero growth rates (4, 11).

For this study, we used an industrially important bacterium *Bifidobacterium breve*, which has been studied for its probiotic capacity and its health-promoting traits (20, 21). This bacterium has been applied in several food products but most importantly in fermented infant formulas due to its capacity to produce metabolites such as galacto-oligosaccharides that are beneficial for infants (22, 23).

The natural environment of *B. breve* is one of the most complex ecosystems that exists: the human gastrointestinal tract (24–26). The human gastrointestinal tract harbors a dense population of microorganisms, which generates a low concentration of nutrients despite its constant inflow. This means that this bacterium often encounters extreme nutrient limitation in the human gut resulting in slow growth (27). *B. breve* has never been studied in a continuous fermentation nor in retentostat systems.

The aim of this research was to investigate the physiological response of *B. breve* NRBB57 at near-zero growth rates using retentostat cultivation with focus on culturability, metabolism and maintenance requirement. Proteome analysis was used to link metabolic and physiological changes to changes in the protein abundances in corresponding metabolic pathways and other relevant functions contributing to the performance of *B. breve* NRBB57 cells at near-zero growth conditions. Implications for *B. breve* ecophysiology and applications are discussed.

## MATERIALS AND METHODS

### Bacterial isolate and storage

In this study, the strain *Bifidobacterium breve* NRBB57 obtained from the strain collection of Danone Nutricia research (Utrecht, The Netherlands) was used. This strain was stored in glycerol stocks (30%) at −80°C until use.

### Culture conditions

Glycerol stocks were streaked on TOS-Propionate agar (Merck, Germany) and incubated at 37°C for 48 hours in anaerobic jars (Advanced Instruments, USA) containing anaerobic gas-generating sachets (Oxoid™ AnaeroGen™). For overnight cultures, single colonies were resuspended in 10 ml TOS-Propionate broth (Merck, Germany), which was made according to manufacturer’s instructions while agar was removed by filtration over a 520B qualitative filter paper (Whatman, United Kingdom) prior to autoclaving the broth filtrate. Overnight cultures were incubated 20 hours at 37°C in anaerobic jars containing anaerobic gas-generating sachets, before they were used to inoculate the chemostat and retentostat cultures.

### *Bifidobacterium breve* culture medium

For the chemostat and retentostat cultivations a partially chemically defined medium was used. This medium contained per kg: 5.4 g lactose.H_2_O as carbon source, 2.7 g KH_2_PO_4_, 0.25 g MgSO_4_.7H_2_0, 10 g bacto-tryptone, 0.5 g L-cysteine-HCl, 1 g yeast extract, 0.002 g D-biotin, 0.0005 g cyanocobalamin, 0.01 g Ca-(D+) panthothenate, 0.005 g nicotinic acid, 0.005 g p-aminobenzoic acid, 0.005 g thiamin-HCl, 0.008 g pyridoxamine-HCl, 0.001 g riboflavin, 0.01 g adenine, 0.01 g xanthine, 0.01 g guanine, 0.01 g uracil, 0.05 g MgCl_2_.6H_2_O, 0.02 g MnSO_4_.H_2_O, 0.005 g ZnSO_4_.7H_2_O, 0.0025 g CuSO_4_.5H_2_O, 0.005 g FeCl_2_.4H_2_O, 0.01 g CaCl_2_.2H_2_0. All the components were properly mixed and the pH was adjusted to 6.5. Finally, the medium was filter-sterilized with a 0.2 μm filter.

### Chemostat cultivation

*B. breve* NRBB57 was grown in chemostats in biological triplicates at dilution rates of 0.12, 0.25 and 0.4 h^−1^ in 0.5 L bioreactors and 0.025 h^−1^ in 1 L bioreactors (Multifors, Infors HT, Switzerland). The temperature was kept constant at 37°C along the fermentation, the pH was controlled at 6.5 by automatic addition of 5 M NaOH and the stirring speed was set at 300 rpm. For anaerobic conditions, the headspace was flushed with gas composed by 5% CO_2_, 5% H_2_, 90% N_2_ at a rate of 0.06 L/min. Bioreactors were inoculated with 1% of an overnight culture made in TOS-Propionate broth (Merck, Germany) as previously described. Bacteria were grown until the end of the exponential phase. Subsequently, the supply of fresh medium was turned on to start the chemostat mode at the dilution rates previously mentioned. Samples were taken once the steady state was reached, which was considered to be achieved after a minimum of 5 volume changes.

### Retentostat cultivation

Biological triplicate retentostats were carried out in 1 L bioreactors (Multifors, Infors HT, Switzerland). Temperature (37°C), pH (6.5), stirring speed and the anaerobic environment were kept constant as previously described for the chemostat cultures. The bioreactors were inoculated with an overnight culture (1%) made in TOS-Propionate broth (Merck, Germany) as previously described. Bacteria were allowed to grow until the end of the exponential phase after which fresh medium supply was turned on to start the chemostat mode at a dilution rate of 0.025 h^−1^. After the chemostat reached the steady state, a polyethersulfone crossflow filter (0.2 μm; Spectrum laboratories, USA) was connected to an outer loop in the effluent line to begin the retentostat mode. Retentostats were run for approximately three weeks. Samples were taken every 2 to 3 days.

### Culturability

To determine the concentration of culturable cells, samples were serially diluted in peptone physiological salt solution (PPS; Tritium Microbiology, The Netherlands) and appropriate dilutions were spread plated on TOS-Propionate agar (Merck, Germany). Plates were incubated anaerobically at 37°C for 48-72 hours, after which the colonies were counted.

### Viability

The culture viability was determined using the LIVE/DEAD *Bac*Light^*TM*^ kit (Molecular Probe Europe, Leiden, The Netherlands) according to the manufacturer’s instructions. The fluorescence of the cells in the treated sample was visualized with a fluorescence microscope (Olympus) at a magnification of 630 times. Bacterial cells with a compromised membrane are considered dead and will stain red with Propidium iodide (PI). Meanwhile cells with an intact membrane were stained green with the DNA probe Syto 9 which are presumed viable. Cells were counted up to 100 per sample and the percentage of viable cells was calculated.

### Cell counts

To determine the cell concentration, samples taken from the bioreactor were diluted 10 to 100 times in peptone physiological salt solution and 25 μl was placed in a cell counting chamber (CellVision technologies, The Netherlands). Subsequently, bacterial cells were counted in a phase contrast microscope (Olympus, Japan) at a magnification of 1000 times. Chains of cells were counted as one, assuming that chains would only form one colony.

### Cell dry weight

Approximately 3 ml of sample (previously weighted) taken from the bioreactor were passed through a pre-weighed 0.2 μm filter (Pall corporation, USA) with the help of a vacuum filtration pump. Filters were then washed with approximately 40 ml of demineralized water and dried at 80°C for 48 hours. Afterwards, the filters were weighed on an analytical balance again to calculate the cell dry weight concentration of the sample. Cell dry weights were measured in duplicate for every sampling point.

### Scanning electron microscopy

The morphology of *B. breve* NRBB57 cells in the retentostat was investigated with scanning electron microscopy (SEM). The samples were collected at time 0 (chemostat) and 7, 15 and 20 days after switching to retentostat mode, centrifuged at 17000×g for 1 minute and the pellets were frozen at −20°C until use. Cells were visualized by SEM as previously described by (11). Briefly, samples were thawed and resuspended in PPS. For every sample, a drop of the suspension was placed in a poly-L-lysine coated coverslip (Corning BioCoat, USA) and left for 1 hour at room temperature. Then, the coverslips were rinsed with phosphate buffered saline, and fixed with 3% glutaraldehyde buffer for 1 hour. Afterwards, the samples were dehydrated in a graded series of ethanol followed by drying with CO_2_ (Leica EM CPD 300, Leica Microsystems, Germany). The coverslips were fitted onto sample stubs with carbon adhesive tabs and sputter coated with 10 nm tungsten (Leica SCD500). Finally, samples were imaged at 2 KV, 6 pA, at room temperature in a field emission scanning electron microscope (Magellan 400, FEI Company, USA).

### Metabolite analysis

Samples were taken directly from the bioreactor, centrifuged at 17000×g for 1 minute and supernatants were stored at −20°C until quantification of extracellular metabolites by High Performance Liquid Chromatography (HPLC) and Ultra-high Performance Liquid Chromatography (UPLC).

For the quantification of lactose, lactate, acetate, ethanol, formate, glycerol and succinate, samples were deproteinated by addition of 100 μl cold Carrez A (0.1 M potassium ferrocyanide trihydrate) to 200 μl sample. After mixing, 100 μl cold Carrez B (0.2 M zinc sulfate heptahydrate) was added, followed by mixing and centrifugation at 17000×*g* for 10 minutes. 10 μl of the deproteinated sample was injected on an Ultimate 3000 (Dionex, Germany) equipped with an Aminex HPX-87H column (300×7.8 mm) with guard-column (Biorad). As mobile phase, 5 mM H_2_SO_4_ was used at a flow of 0.6 ml/min. The column temperature was kept at 40°C. Compounds were detected by refractive index detector (RefractoMax 520). The analysis was performed with technical duplicates.

Amino acids (alanine, asparagine, aspartic acid, cystine, glutamic acid, glutamine, arginine, glycine, histidine, isoleucine, leucine, lysine, methionine, phenylalanine, proline, serine, threonine, tryptophan, tyrosine, valine) and ammonia (NH_3_) were quantified by UPLC. 40 μl supernatant was deproteinated by addition of 50 μl 0.1 M HCl, containing 250 μM norvaline as internal standard and 10 μl 30% sulfosalicylic acid (SSA). Subsequently, the solution was mixed and centrifuged at 17000×*g* for 10 minutes at 4 °C. Amino acids and ammonium were derivatized using the AccQ•Tag Ultra derivatization kit (Waters, USA). 20 μL of the deproteinated supernatant or standard amino acids mixture was mixed with 60 μL AccQ•Tag Ultra borate buffer in glass vials. For deproteinated samples 75 μL of 4 M NaOH was added to 5 ml borate buffer to neutralize the addition of SSA. Subsequently, 20 μL of a AccQ•Tag reagent dissolved in 2.0 mL AccQ•Tag Ultra reagent diluent was added and immediately vortexed for 10 seconds. Then, the sample solution was capped and heated at 55 °C in a heatblock for 10 minutes. Amino acids and ammonium were quantified by UPLC by injection of 1 μl sample on an Ultimate 3000 (Dionex, Germany) equipped with a AccQ•Tag Ultra BEH C18 column (150 mm x 2.1 mm, 1.7 μm) (Waters, USA) with BEH C18 guard column (5 mm x 2.1 mm, 1.7 μm) (Waters, USA). The column temperature was set at 55°C and the mobile phase flow rate was maintained at 0.7 ml/min. Eluent A was 5% AccQ•Tag Ultra concentrate solvent A and Eluent B was 100% AccQ•Tag Ultra solvent B. The separation gradient was 0-0.04 min 99.9% A, 5.24 min 90.9% A, 7.24 min 78.8% A, 8.54 min 57.8% A, 8.55-10.14 min 10% A, 10.23-17 min 99.9% A. Compounds were detected by UV measurement at 260 nm. Glutamine and arginine could not be separated in the UPLC analysis.

### Modeling biomass accumulation and estimation maintenance

The accumulation of biomass in the retentostat cultivations were modelled using a modified Verseveld equation in which metabolic changes are taking into account (11) (Eq. 1). Moreover, we assumed that the maintenance coefficient gradually decreases towards near-zero growth rates as found for *Lactococcus lactis* (Eq. 2), (11), in which the growth rate was calculated by taking the first derivative of Eq. 1 giving Eq. 3.

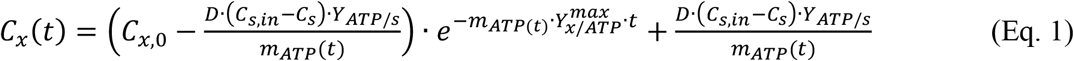

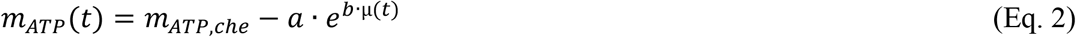

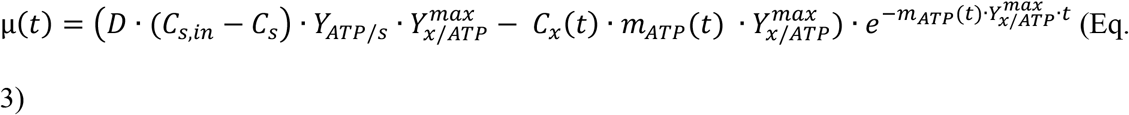

in which t is the time (h), C_x_ is the biomass concentration (gDW/kg), C_x,0_ is the biomass concentration at *t* = *t* − *Δt* (gDW/kg), D is the dilution rate (h^−1^), C_s,in_ is the substrate concentration in the medium (15 mmol/kg lactose), C_S_ is the substrate concentration in the effluent, Y_ATP/s_ is the ATP yield on substrate (mol ATP/CmolS), m_ATP_ is the maintenance coefficient (mol ATP.gDW^−1^.h^−1^), Y_x/ATP_^max^ is the maximum biomass yield on ATP measured in chemostat cultivation (gDW/mol ATP), m_ATP,che_ is the maintenance coefficient measured in chemostat cultivations (mol ATP.gDW^−1^.h^−1^), μ is the growth rate (h^−1^), a and b are parameters describing the relation between the maintenance coefficient and the growth rate.

The Y_ATP/s_ was calculated based on the measured metabolite production (Eq. 4) assuming lactose was the only energy source, 1 mol ATP is produced per mol acetate produced in the bifid shunt, while 2 mol ATP is produced per mol acetate produced via pyruvate formate lyase and 1 mol ATP is produced per mol lactate and ethanol:

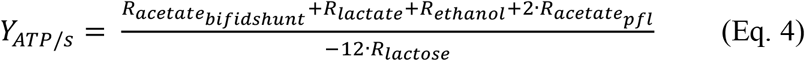

in which R_i_ is the production rate (mol/h) of compound i. Because in the bifid shunt 1 mol lactose is converted to 3 mol acetate, equation 4 can be rewritten as:

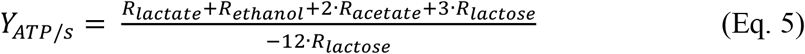

Input data for the modelling were online optical density measurements, which were converted to biomass dry weight concentrations using a second-order polynomial relation. To fit the model, the variable parameters a and b were optimized by minimizing the sum of squared errors between the model and the data in 10-minute time intervals. This was done using the solver add-in of Excel.

### Proteome analysis

For the proteome analysis, samples were taken from the steady states of 3 biological replicates of the chemostats (μ=0.4 h^−1^, μ=0.12 h^−1^, μ=0.05 h^−1^, μ=0.025 h^−1^) and from 3 biological replicates of the retentostats after 1, 2 and 3 weeks after connecting the filter. Samples were centrifuged at 13000×*g* for 5 minutes and pellets were frozen at −80°C until use.

Cell lysis was achieved through bead beating in a FastPrep-24TM 5G instrument (MP Biomedicals) for 6 times 30 seconds at 6.5 m/s with cooling after every bead step. Protein concentration was assessed using Pierce BCA protein assay (ThermoFisher Scientific, Waltham, USA; S7). Subsequent protein digestion was performed overnight using dithiothreitol (DTT, 2 mM), iodoacetamide (IAA, 4 mM) and trypsin (1:50 of a 1 μg/μL solution) at 37°C. Cleanup was performed through SPE columns (Solid Phase Extraction; ThermoFisher Scientific, Waltham, USA) with acetic acid (100 mM in 95% acetonitrile) to be finally dissolved in the eluent acetic acid (100 mM). Samples were analyzed by Nano liquid chromatography-high-resolution mass spectrometry (nano-LC-HRMS/MS) as previously described (28). An Agilent 1200 HPLC system (Agilent Technologies, Santa Clara, USA) was used, connected to a Q-Exactive Plus Mass spectrometer (ThermoFisher Scientific, Waltham, USA). Peptides were trapped on a 100 μm inner diameter trap column packed using ReproSil-Pur C18-AQ, 3 μm resin (Dr. Maisch, Ammerbuch, Germany) at 5 μL/min in 100 mM acetic acid. Afterwards the peptides were eluted at 100 nL/min in a 90 minutes extended gradient from 10-40% acetic acid solvent (in 95% acetonitrile) to a 20-cm IntegraFrit column (50 μm inner diameter, Reprosil-Pur C18-AQ 3 μm, New Objective, Woburn, USA). The acquired spectra were analyzed using Thermo Proteome Discoverer in combination with Mascot (ThermoFisher Scientific, Waltham, USA). The reference database comprised of protein sequences from *B. breve* NRBB57 from Uniprot and typical contaminants. Relative protein quantification was performed by Proteome Discoverer based on peptide intensity signals using default settings. The obtained abundances of all detected proteins are listed in Table S1.

### Proteome data analysis

To determine significant differences in the proteome between growth rates, the abundances of the proteins were plotted vs the growth rate. Afterwards, linear regression was calculated to find the slopes of the line and check how significant this slope differed from 0 (p-value) in R (v3.63). The significantly different abundances were considered when the p-value of the slope was <0.05. Finally, proteins that significantly changed were clustered with GSEA-pro v3.0 software (http://gseapro.molgenrug.nl/). The proteins were clasified into clusters of Orthologous Groups (COG) to predict the functions of the different sets of proteins.

## RESULTS

In this study, *Bifidobacterium breve* NRBB57 was anaerobically cultivated for 3 weeks in three independent retentostat cultivations in a medium containing lactose as growth-limiting substrate. During cultivation, several parameters were measured aiming to establish the physiological adaptations of this strain to near-zero growth rates. One of the replicates was stopped after 2 weeks due to clogging of the cross-flow filter and is therefore only included in the proteome analysis.

### Biomass accumulation

In the retentostat cultivations, the biomass concentration gradually increased 6-fold (Fig. 1A), while the growth rate decreased progressively to 0.00092±0.00006 h^−1^ (Fig. 1B).

**Figure 1:**
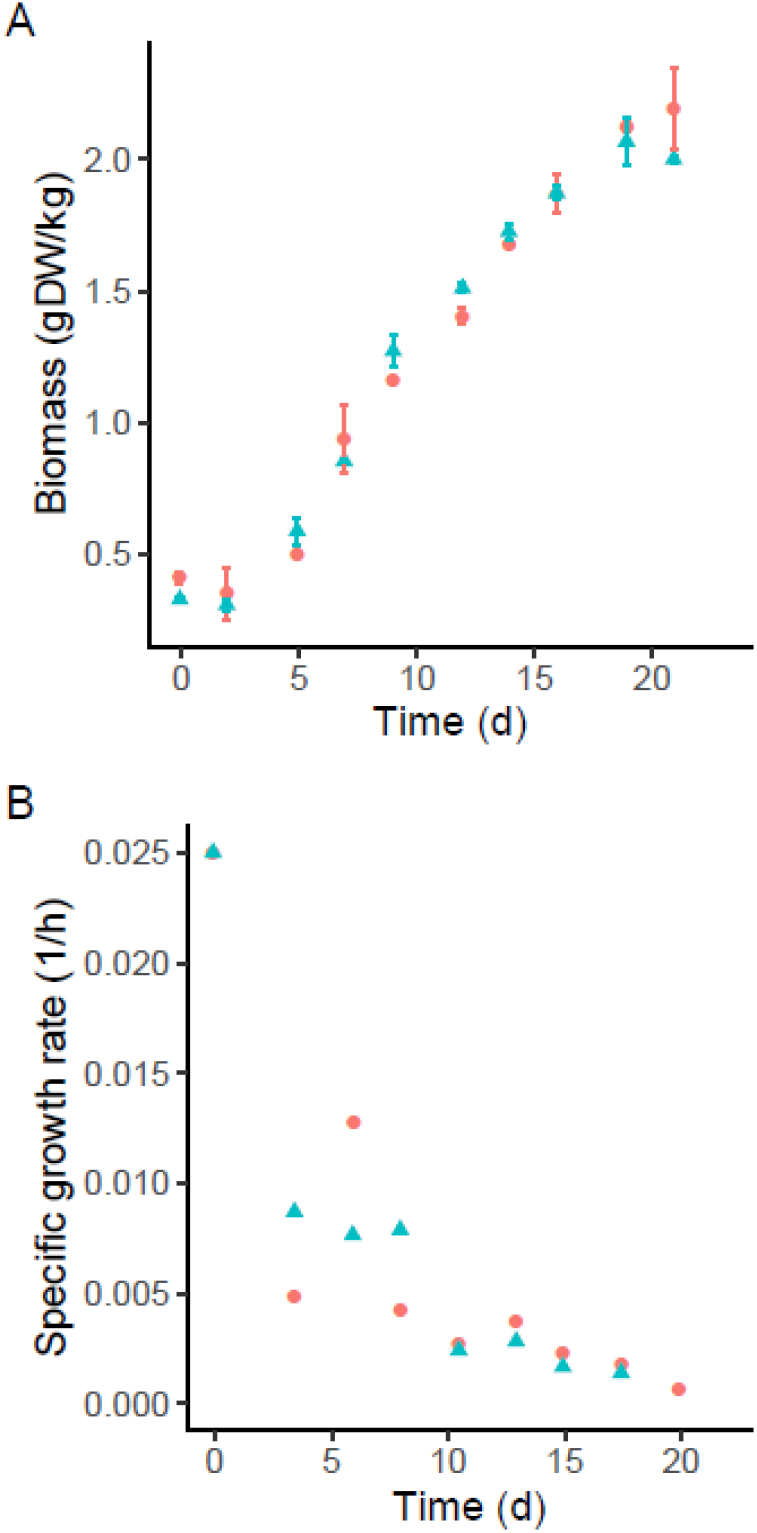
Growth of *B. breve* NRBB57 during the retentostat cultivations. Colors and symbols represent different biological replicates. Time zero corresponds to the last day of the chemostat at a growth rate of 0.025 h^−1^, when retentostat cultivation mode was started by the connection of a filter in the effluent line. (A) Biomass accumulation throughout the retentostat cultivation. Data points corresponds to the average ± standard deviation of technical duplicates. (B) Calculated specific growth rates based on cell dry weight measurements.

### Culturability, viability and morphology

Culturability was estimated by plating serial dilutions on TOS-Propionate agar plates and comparing the number of culturable cells with the total number of cells estimated using a counting chamber (chains of cells were counted as 1). Colony Forming Units (CFU/ml) decreased over time (Fig. 2A), while the cell counts increased approximately 10-fold (Fig. 2B).

**Figure 2:**
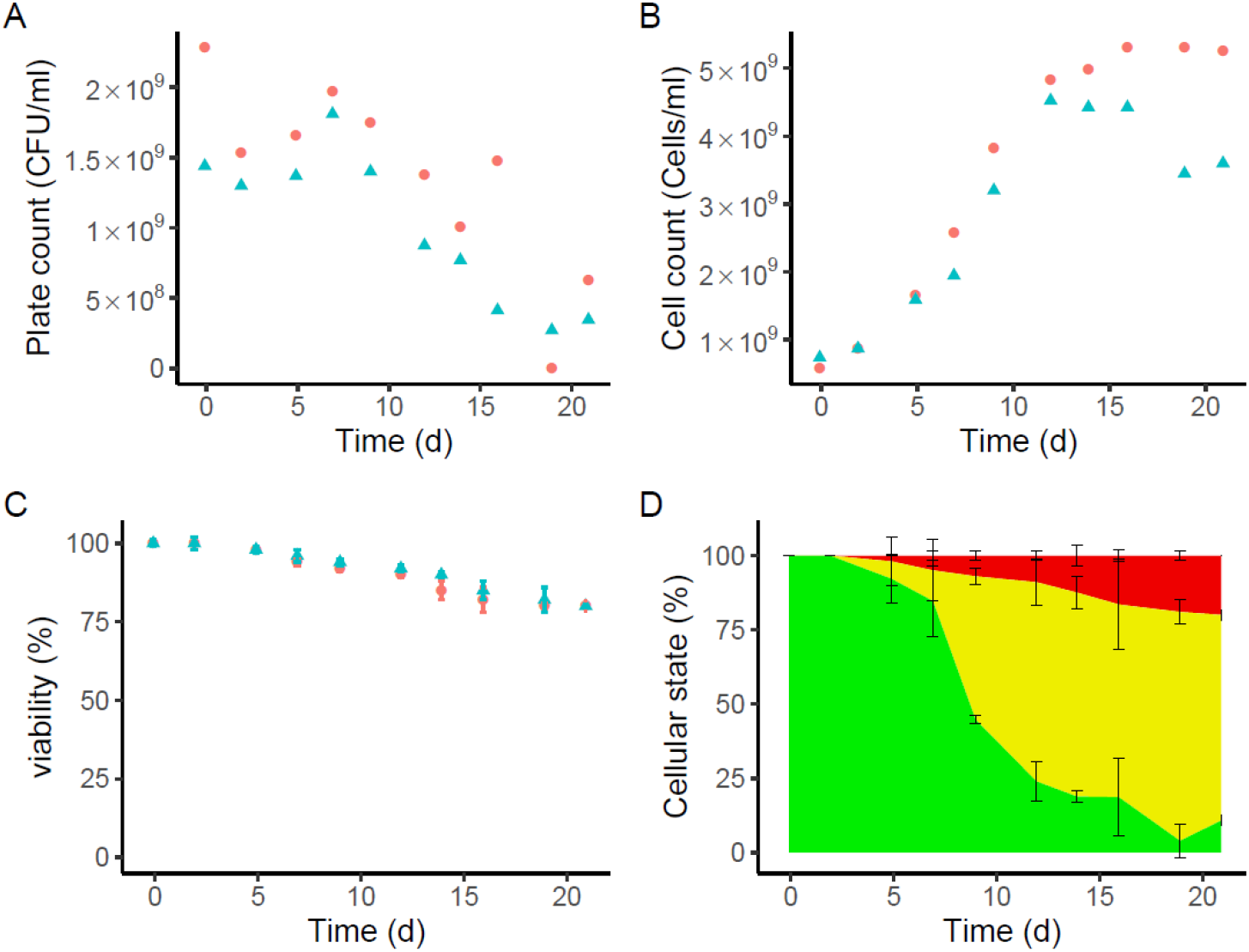
Quantification of cell counts, plate counts, viable cells and cellular state of *Bifidobacterium breve* NRBB57 during retentostat cultivation. Colors and symbols represent different biological replicates. Time zero corresponds to the last day of the chemostat at a growth rate of 0.025 h^−1^, when retentostat cultivation mode was started by connecting a filter to the effluent line. (A) Culturable cells were estimated by plate counts. Every data point corresponds to the average of a technical duplicate. (B) Cell counts measurements during the retentostat cultivation. Every time point corresponds to the average of a technical replicate. Total number of cells were counted using a counting chamber. Chains of cells were counted as 1. (C) Viability during the retentostat cultivation assessed by live/dead staining. (D) Physiological status of the cells calculated based on data from panel A, B and C: green, yellow and red, represent fractions of culturable cells, viable-but-non-culturable cells and dead cells, respectively.

Likewise, we determined the viability of the culture by assessing the membrane integrity of the cells using the LIVE/DEAD *Backlight*™ kit. Bacterial cells were classified into two categories: red PI-stained cells conceivably had damaged membranes and therefore were considered not viable, while green SYTO 9-stained cells had intact membranes and were considered viable. The viability decreased from 99% in chemostat mode (t=0) to 80% at the end of the retentostat culture (Fig. 2C). These observations indicate that most of the bacterial population in this culture stayed viable during the three weeks of retentostat cultivation, but lost the ability to grow on TOS-Propionate agar plates (approximately 70%) (Fig. 2D).

The morphology of the *B. breve* NRBB57 was assessed by scanning electron microscopy (SEM). This showed that the morphology changed in the first and second week of retentostat cultivation when cells displayed corrugation (Fig. 3). By the third week, bacterial cells became much longer and showed branching which were counted as 1 as mentioned above.

**Figure 3:**
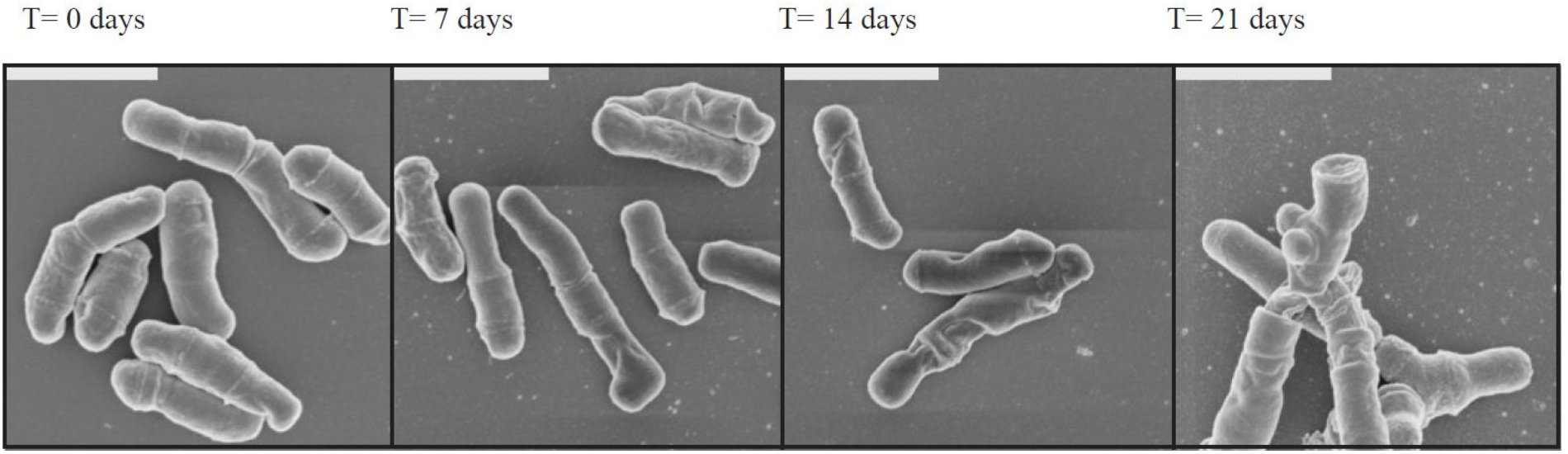
Morphology of *B. breve* NRBB57 cultivated in the retentostat and visualised under scanning electron microscopy. Samples were taken at time 0 which corresponds to the last day of the chemostat at a growth rate of 0.025 h^−1^ and after 7, 14 and 21 days of retentostat cultivation. The length of the white bars in the top left corner of the photographs correspond to 1 μm.

### Metabolism

Concentrations of the products of lactose metabolism (acetate, lactate, ethanol, formate) were measured along with the production of succinate and the consumption of glycerol (Fig. 4). At the start of the retentostat cultivation, lactose was mainly converted into acetate, ethanol and formate indicating that *B. breve* NRBB57 metabolized lactose via the bifid shunt (29). This behaviour was also observed in chemostat cultivations of *B. breve* NRBB57 at higher growth rates up to 0.4 h^−1^, while in batch cultures this strain produces a mixture of acetate and lactate (Fig. S1). Throughout the retentostat cultivation, acetate production was stable while the production of ethanol and formate slightly decreased, and lactate production increased indicating a small but gradual change in pyruvate dissipation while the growth rate decreases. Furthermore, the concentration of glycerol decreased starting after approximately 1 week of retentostat cultivation, indicating that glycerol was consumed. Interestingly, the concentration of succinate increased over time, which could point to catabolism of amino acids. Therefore, we decided to also assess the concentration of amino acids during the fermentations.

**Figure 4:**
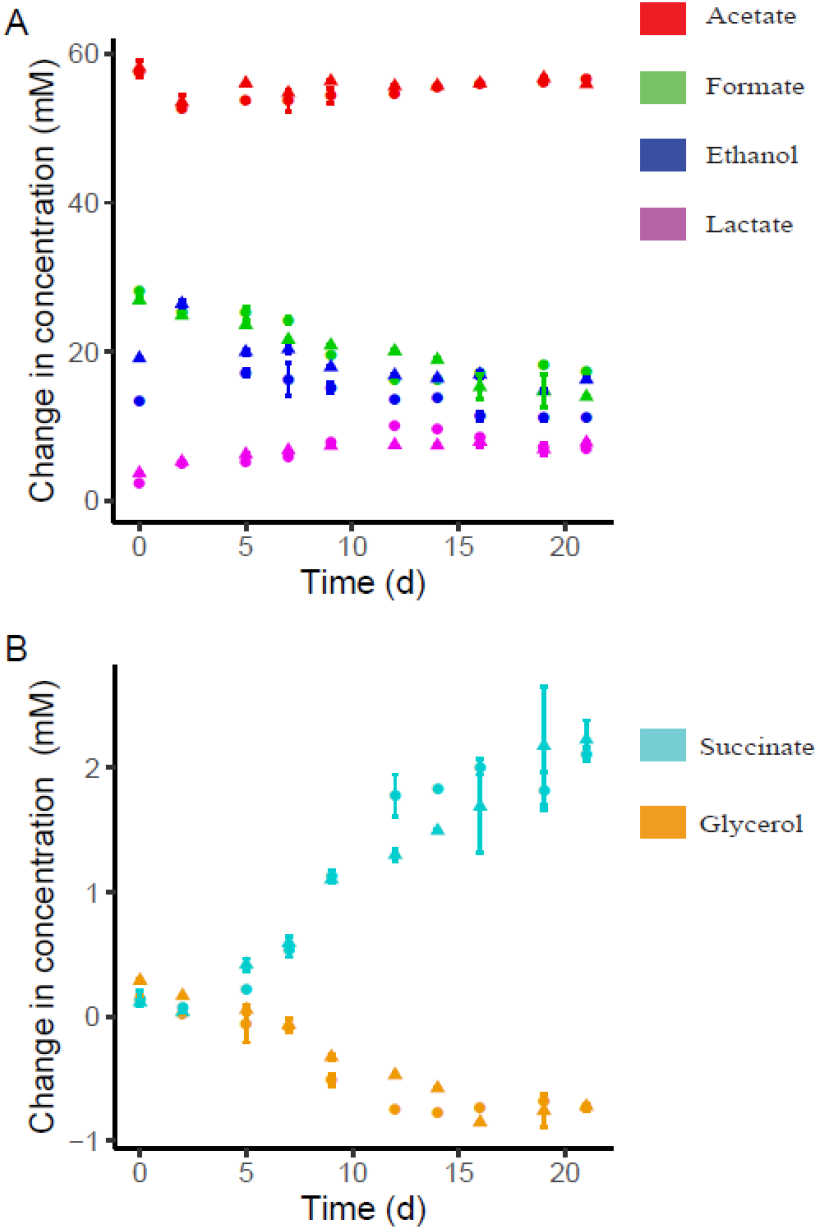
Extracellular metabolite concentrations in the retentostat over time. Colors represent different metabolites and symbols represent different biological replicates. Data points corresponds to the average ± standard deviation of technical duplicates. (A) Main products of the lactose metabolism. (B) Secondary metabolism of succinate and glycerol. Negative production for glycerol indicates consumption.

### Amino acids

Concentrations of amino acids were measured by UPLC. During the retentostat cultivation the concentration of several amino acids decreased (aspartic acid, glutamine/arginine, serine) (Fig. 5). While the succinate concentration gradually increased 2 mM from time 0 until the end of the retentostat cultivation, the aspartic acid and glutamine/arginine concentration decreased by only 0.1 mM and 0.75 mM, respectively. This suggests that amino acid metabolism could only partially explain the increase in succinate indicating that the extra succinate originates from another source. Likewise, the ornithine concentration increased 0.15 mM conceivably as the end product of arginine catabolism. Ammonia (NH_3_) production also increased during the retentostat cultivation (1 mM), which correlates with the increased utilization of amino acids. Concentrations of other amino acids (alanine, asparagine, cystine, glutamic acid, glycine, histidine, isoleucine, leucine, lysine, methionine, phenylalanine, proline, threonine, tryptophan, tyrosine, valine) did not decrease suggesting that these were not utilized (Fig. 2S).

**Figure 5:**
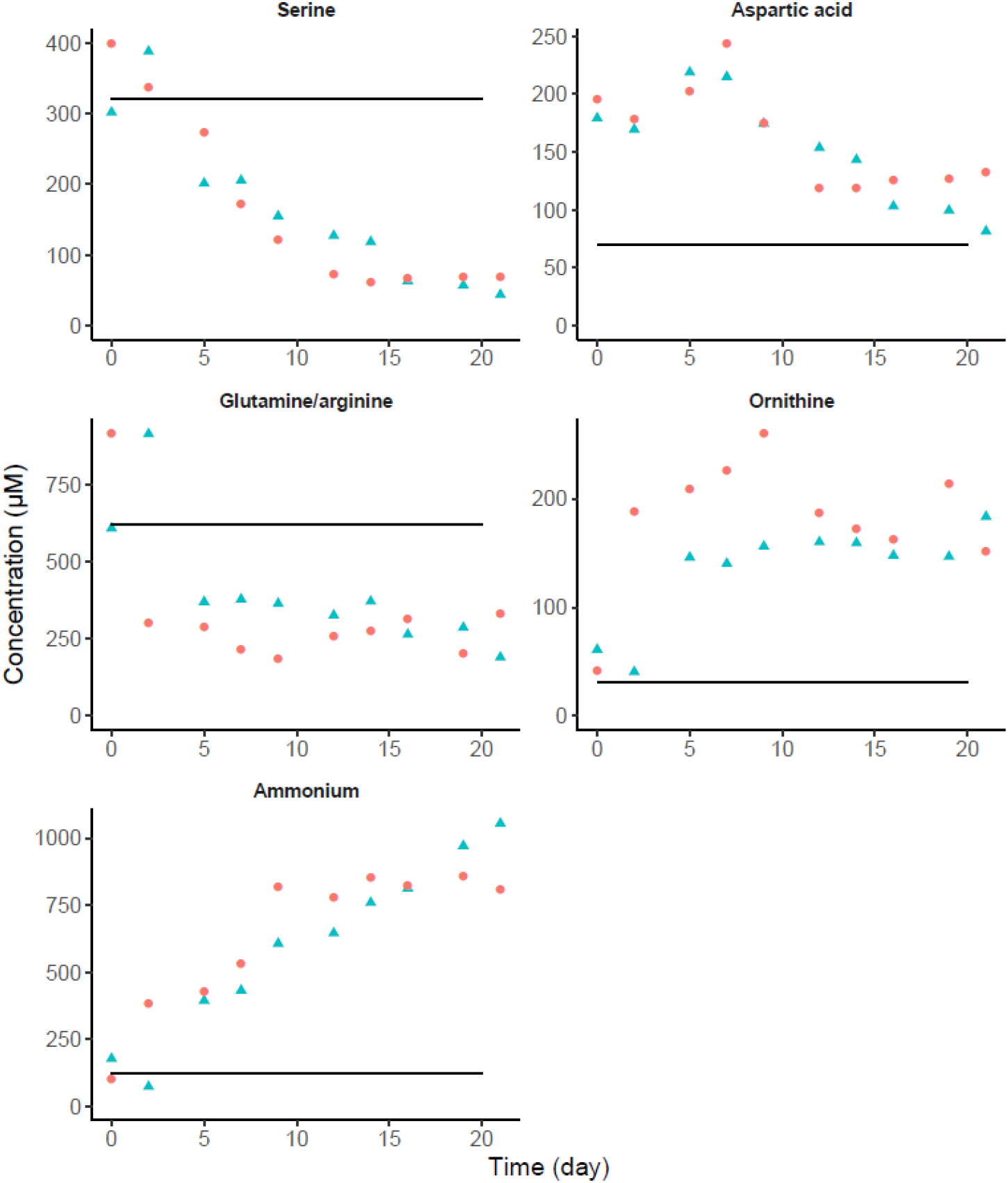
Extracellular amino acids concentrations and ammonia in the retentostat measured with UPLC over time. Colors and symbols represent different biological duplicates. Time zero represents the last day of the chemostat mode. The horizontal line represents the concentration of the amino acids and ammonia in unused media. Glutamine and arginine could not be separated in the UPLC and the decreased concentration could be due to either increased glutamine and/or arginine utilization. Only a selection of the amino acids is shown. All quantified amino acids can be found in (Fig. S2).

### Maintenance requirement

Metabolite analysis suggests that *B. breve* NRBB57 used alternative energy sources under the extremely energy-limited conditions, most likely to produce as much energy as possible thereby improving its chance of survival. Therefore, we investigated if and to what extend *B. breve* NRBB57 is capable of decreasing the energy required for its maintenance purposes. To estimate the maintenance requirements at near-zero growth rates, biomass accumulation in the retentostat cultures has been modelled using a modified Verseveld equation, in which ATP production was estimated based on the metabolite production (11) as explained in the experimental procedures. The maximum biomass yield on ATP (Y_x/ATP_^max^) and maintenance coefficient (m_ATP,che_) were determined in chemostat cultivations at dilution rates between 0.025 and 0.4 h^−1^ and estimated to be 16.73±0.85 gDW/mol ATP (estimate ± standard error) and 3.32±0.77 mmol ATP.gDW^−1^.h^−1^ (estimate ± standard error), respectively (Fig. 6A). Comparing the m_ATP,che_ with the biomass specific ATP production rate (q_ATP_) during the retentostat cultivation showed that the maintenance coefficient reduced at least 3.6-fold. Therefore, we assumed in the modelling that the maintenance coefficient gradually decreases towards near-zero growth rates as found for *Lactococcus lactis* (11). Biomass accumulated in the cross-flow filter in the first couple of days after connecting the filter resulting in significant underestimation of the biomass production. Therefore, biomass accumulation in the bioreactor was modelled from 5 days onwards resulting in a good model fit (square root of mean squared error is 0.066 and 0.084 gDW/kg (Fig. S3). The model estimated that during 3 weeks of retentostat cultivation the maintenance coefficient reduced 3.6-fold to 0.915±0.016 mmol ATP.gDW^−1^.h^−1^ (Fig. S4). The estimated growth rates after 1, 2 and 3 weeks of retentostat cultivation were 0.0055±0.0011, 0.0018±0.00005 and 0.00092±0.00006 h^−1^, respectively (Fig. S5). Finally, the estimated growth rates in the retentostat cultures were plotted against the biomass specific ATP production rate and compared with data from the chemostat cultures used to estimate the Y_x/ATP_^max^ and m_ATP_ (Fig. 6B). This showed that these estimates are only valid at growth rates above 0.025 h^−1^ because at near-zero growth rates the ATP required for maintenance purposes is significantly reduced.

**Figure 6:**
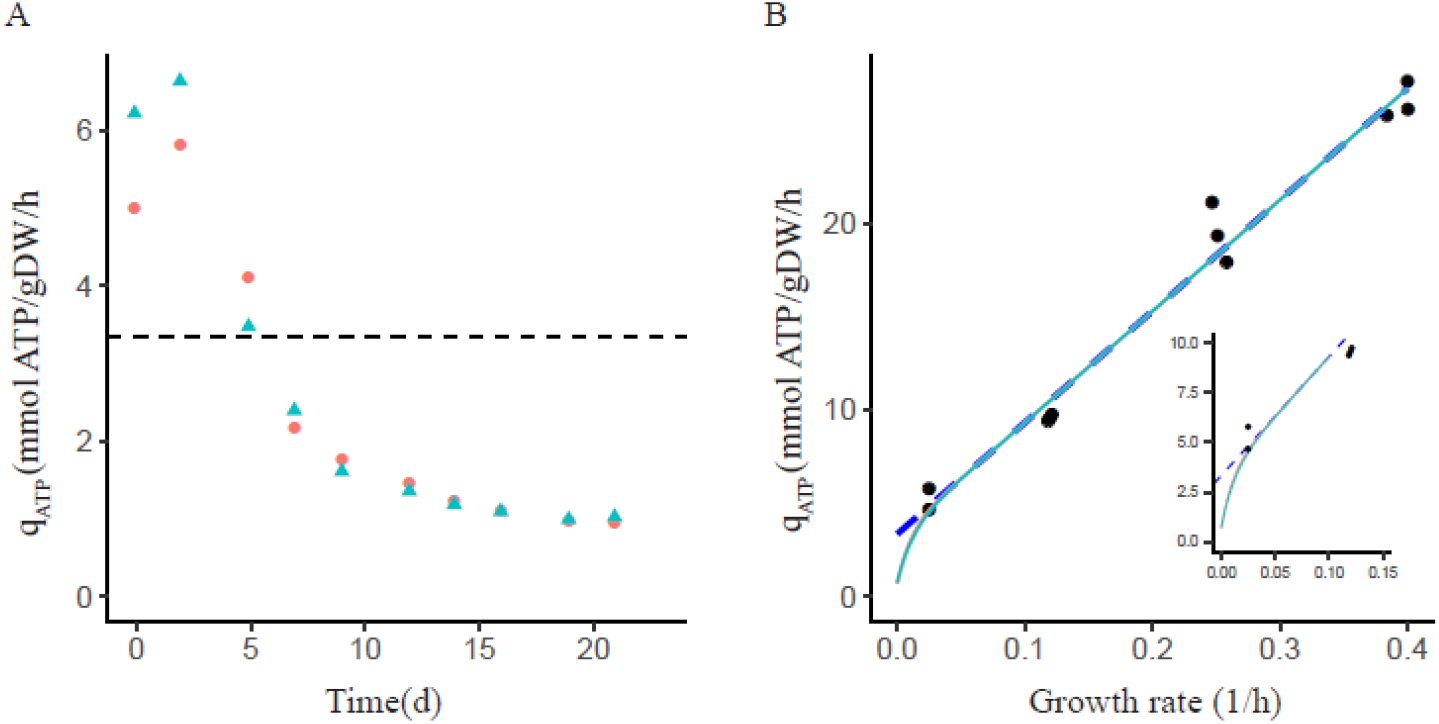
Model predictions on the biomass accumulation, growth rate and the biomass specific ATP production rate during retentostat cultivations. Colors and symbols represent different biological duplicates. (A) Biomass specific ATP production rate (q_ATP_). The black dashed line indicates the m_ATP_ estimation from the chemostat. (B) Relationship between the growth rate and the biomass specific ATP production rate during chemostat and retentostat cultivations. Black dots represent the data from chemostat cultures. The blue dashed line represents the conventional prediction of the relationship based on only chemostat cultures. The solid lines represent the linear regression used for estimation of the biomass yield on ATP. The solid lines also take into account the retentostat cultures for this relationship assuming a decreasing maintenance requirement at near-zero growth rates. Axis titles in the inset are the same as in the main figure.

### Proteome Analysis

The proteome of *B. breve* NRBB57 was analyzed to investigate how this bacterium manages to adapt to the nutrient limitation and consequently decreasing growth rates. Additionally, through this analysis, it was possible to link the phenotypes described in the results above to changes in protein abundances. Samples were taken in biological triplicates from a wide range of growth rates in chemostat cultures (0.4, 0.25, 0,12 and 0.025 h^−1^) and retentostat cultures (0.0055, 0.0018, 0.00092 h^−1^). Protein abundances in all conditions were compared with the chemostat at 0.025 h^−1^, because this condition also served as starting point for the retentostat cultivations. Cluster analysis of all proteome profiles showed that retentostat and chemostat samples clustered separately, but within the group of retentostat samples (μ≤0.0055 h^−1^) they cluster per replicate and not per growth rate (Fig. S6).

To detect proteins with a significantly different abundance as function of the growth rate, the protein abundance ratio 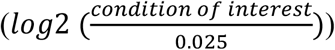 was plotted against the growth rate (on log10 scale). Linear regression was applied on either the high growth rates (μ ≥ 0.025 h^−1^) or near-zero growth rates (μ ≤ 0.025 h^−1^) to find the slope and its significance (p-value). Because all data were expressed as log2 values of the protein abundance ratio compared to values obtained at 0.025 h^−1^, all regression lines were forced to the point (0.025; 0). Subsequently, each protein was allocated to a group based on their slopes and p-values. In total 1134 proteins were detected and 655 out of these showed significant changes in abundance in comparison with the cells cultivated at 0.025 h^−1^ (chemostat cells). Out of these 655 proteins, 249 proteins showed significant changes in abundance in retentostat cells in comparison to chemostat cells. Whereas 78 proteins overlapped between the retentostat and the chemostat (Fig. 7).

**Figure 7:**
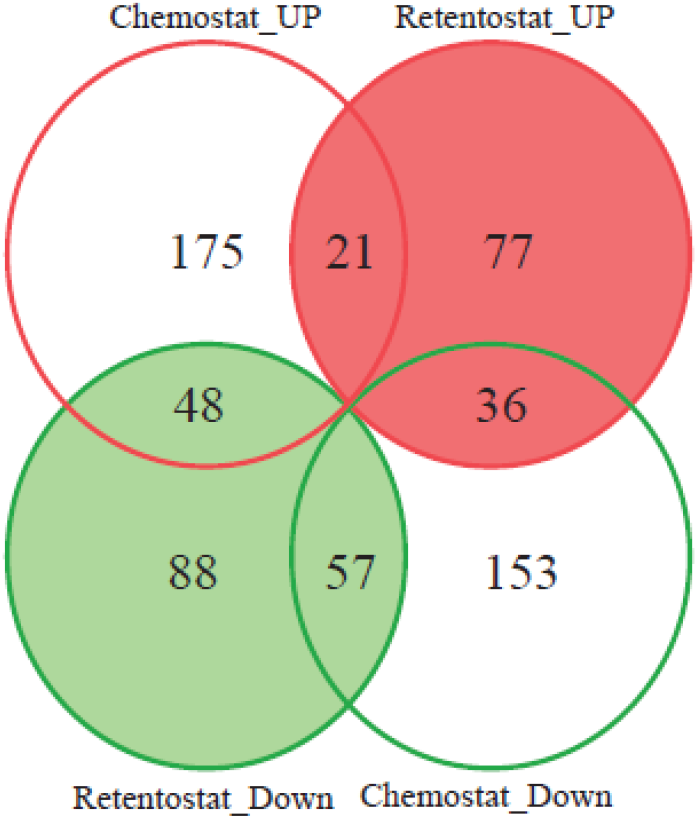
Venn diagram of the proteins with significant changes in their abundance during the chemostat and the retentostat cultivations. Chemostat_UP and Retentostat_UP represent proteins with significant higher abundance at higher growth rates in the range μ≥0.025 h^−1^ and μ≤0.025 h^−1^, respectively. Chemostat_DOWN and Retentostat_DOWN represent proteins with a significant lower abundance at higher growth rates in the range μ≥0.025 h^−1^ and μ≤0.025 h^−1^, respectively. 1135 proteins were identified in total. An overview of the proteins in each group is given in Table S1.

A functional enrichment analysis was performed for proteins that showed significant differences in their abundance in cells cultivated at lower growth rates (retentostat cells) in comparison to their abundance in cells cultivated at 0.025 h^−1^ (chemostat cells). The results show the clusters of orthologus groups (COG) categories to which the proteins with increased and reduced abundances belong to (Fig. 8). Interestingly, among the proteins with increased abundances in the retentostat, the most over-represented categories (highest score assigned by the software) correspond to (i) posttranscriptional modification, protein turnover and chaperones; (ii) translation, ribosomal structure and biogenesis; (iii) nucleotide transport and metabolism; (iv) amino acid transport and metabolism; and (v) replication, recombination and repair. The category related to “translation and replication” includes many proteins that participate in the stress response, such as chaperones and DNA repair proteins. The COG category “nucleotide transport and metabolism” includes proteins that are likely to be involved in (pp)pGpp synthesis, indicating that the severe nutrient limitation in the retentostat induced a stringent response in *B. breve* NRBB57. The category amino acid transport and metabolism consist of proteins involved in the biosynthesis of several amino acids, but also contains several proteins involved in catabolism of serine (serine dehydratase), aspartic acid (aspartate aminotransferase) and arginine (argininosuccinate synthase, ornithine carbamoyltransferase). Finally, other proteins that were more abundant at near-zero growth rates were proteins involved in polyphosphate synthesis (polyphosphate kinase and proteins involved in phosphate transport) as well as several tRNA synthetases.

**Figure 8:**
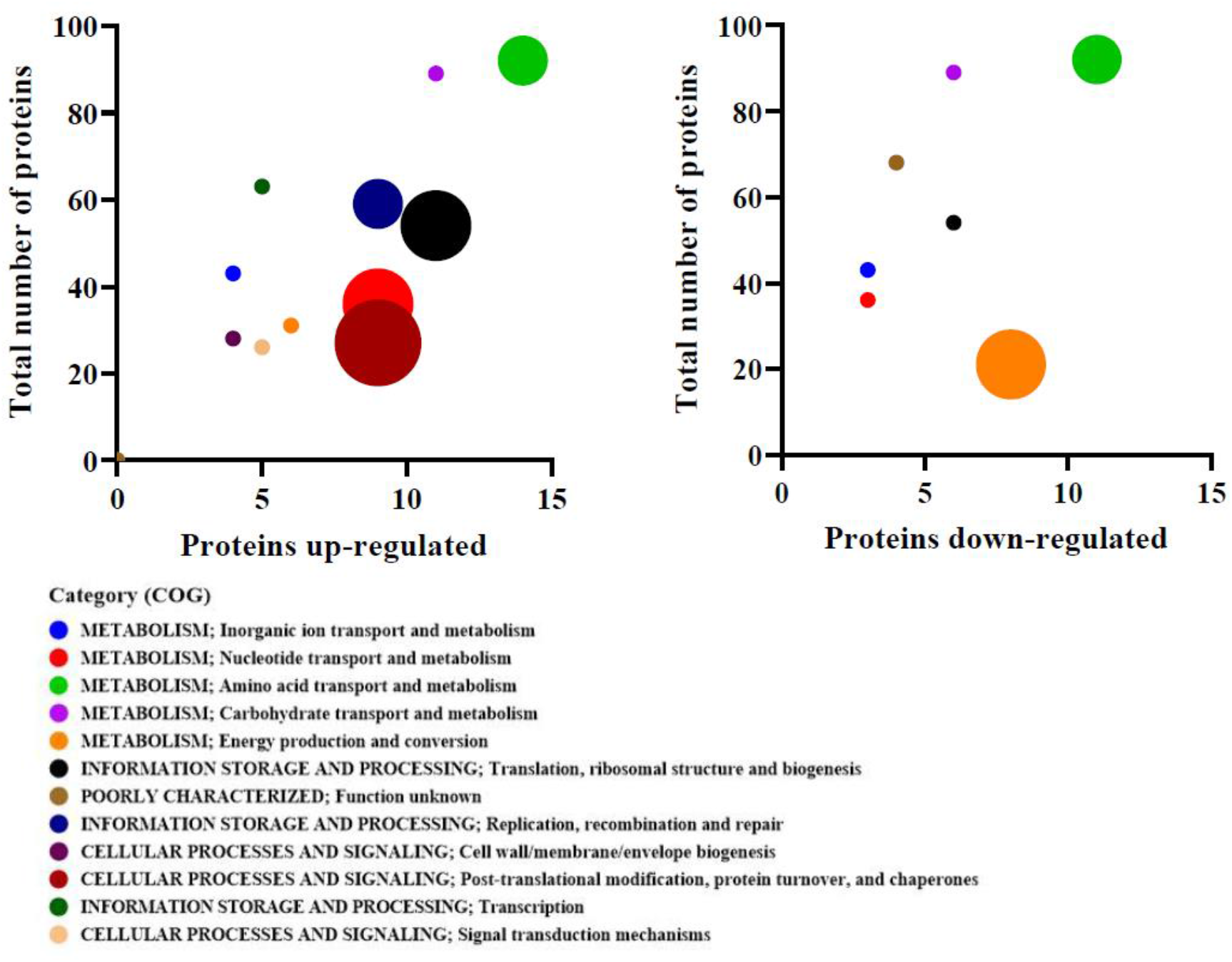
Enrichment of proteins with significant differences in the abundances during the retentostat cultivation. This analysis was carried out with the GSEA-pro v3.0 software. Different colors represent different functional categories. The total number of proteins indicates all the proteins that compose that category versus the proteins that were detected with significant differences. Bubble size represents the score or significance assigned by the software being 3 the biggest size and 0 the smallest size, which indicates that higher scores have more up or down-regulated proteins in relation with the total of proteins in that category.

On the other hand, for the proteins with decreased abundances at near-zero growth rates the most over-represented categories included: energy production and conversion; and amino acid transport and metabolism. Other proteins that were significantly less abundant at near-zero growth rates were ribosomal proteins (15 out of 52) which indicates a reduction in ribosome synthesis.

### Metabolic energy production

Our results show that during the retentostat cultivation, *B. breve* NRBB57 is able to adapt its metabolism to cope with nutrient scarcity. Lactose was mainly converted to acetate and at decreasing growth rates lactate concentrations gradually increased. Simultaneously, formate and ethanol concentrations slightly decreased with increasing residence time in the retentostat bioreactor. Aditionally, the extracellular concentrations of amino acids (serine, aspartic acid and glutamine/arginine) and glycerol were reduced over time in the retentostat. These observations were linked to the proteome data showing that several proteins involved in the catabolism of amino acids, glycerol and lactate production had significantly higher abundances in comparison to the chemostat cultivation. This led to the hypothesis that *B. breve* NRBB57 started using these amino acids as well as glycerol as additional energy sources during declining growth rates caused by nutrient limitation.

Based on the metabolic and proteome analysis, we propose an integration of the catabolic pathways for lactose, arginine, serine, aspartic acid and glycerol and how the utilization of these substrates potentially contribute to metabolic energy generation (Fig. 9). Lactose metabolism by *B. breve* leads to the production of acetate, ethanol, formate and lactate. Moreover, we observed utilisation of arginine concomitantly with the production of ornithine. Simultaneously, serine was probably converted to pyruvate and ammonia. In our model, glycerol is converted to pyruvate and aspartate is proposed to be converted to succinate.

**Figure 9:**
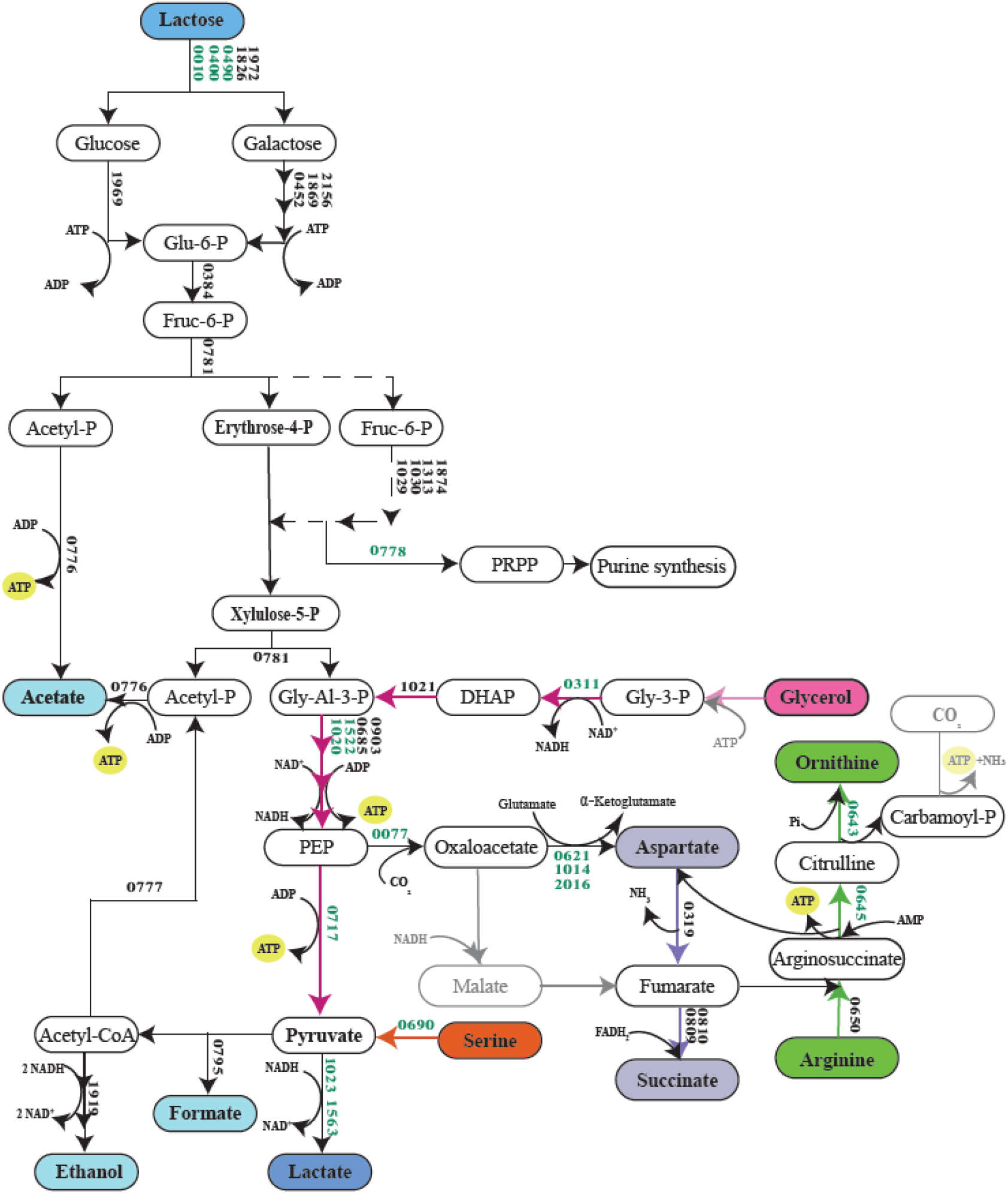
Proposed metabolic energy generating pathways activated in *B. breve* NRBB57 at near-zero growth conditions in the retentostat. Proteins with significantly higher abundances in the retentostat cultivations are shown in green. Reactions not identified in bifidobacteria are shown in grey. In this pathway, lactose is mainly converted to lactate and acetate (2:3). Glycerol is converted to ethanol and serine is converted to acetate. Moreover, aspartate is transformed to succinate and arginine is converted into ornithine. The metabolism of one molecule of lactose delivers 5 ATPs. From serine to acetate 1 ATP is produced. From aspartate to succinate, 1 FADH2 is consumed, which means that 0.5 Acetyl-CoA can be converted to acetate instead of ethanol, which is equivalent to 0.5 ATP. From arginine to ornithine 2 ATPs are produced on the known steps in *B. breve*. Finally the conversion of glycerol to ethanol gives 1 ATP. Abbreviations correspond to: glucose-6-phosphate (Glu-6-P); fructose-6-phosphate (Fruc-6-P); Glyceraldehyde-3-phosphate (Gly-Al-3-P); phosphoribosyl pyrophosphate (PRPP); 2-phosphoenolpyruvate (PEP); dihydroxyacetone phosphate (DHAP); glycerol-3-phosphate (Gly-3-P).

### (p)ppGpp induced stringent response

Enzymes involved in the production of the alarmones guanosine tetraphosphate (ppGpp) and guanosine pentaphosphate (pppGpp) and guanosine 5′-monophosphate 3′-diphosphate (pGpp) (collectively referred as ppGpp were more abundant at near-zero growth rates (Fig. 10). Proteomic data suggested that synthesis of guanosine diphosphate (GDP) and guanosine triphosphate (GTP), which are precursors of ppGpp and (pp)pGpp, respectively, increased at higher growth rates, but the enzymatic complex RelA/SpoT that converts GDP/GTP into (pp)pGpp was less abundant at higher growth rates, which indicates that GDP/GTP was produced solely for growth purposes. In contrast, proteins involved in GTP synthesis also increased in the retentostat cultivation, but now the enzymatic complex RelA/SpoT increased concomitantly indicating that at near-zero growth rates the GDP and the GTP is converted to (pp)pGpp to activate the stringent response.

**Figure 10:**
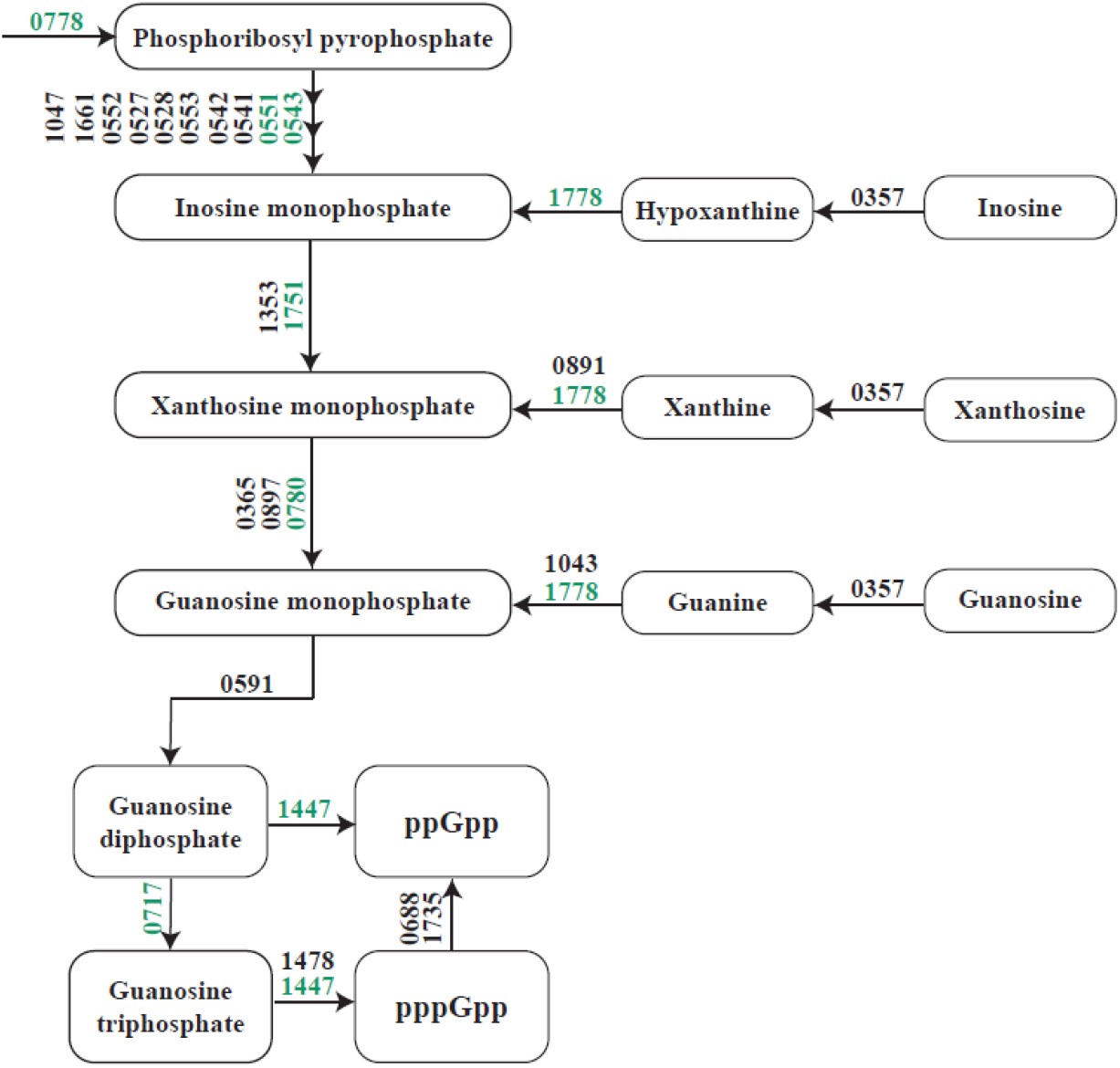
Schematic representation of (p)ppGpp production by *B. breve* NRBB57 in the retentostat cultivation. Proteins with significantly higher abundances in the retentostat cultivations are shown in green.

## DISCUSSION

In the present study, the physiology of *Bifidobacterium breve* NRBB57 has been investigated at near-zero growth rates using retentostat cultivations. In a retentostat cultivation bacterial cells are calorically restricted but not starving. Several parameters were measured in this study, including biomass accumulation, culturability, viability, morphology, metabolite production and consumption, and proteomic adaptations. Retentostat cultivation was carried out for 21 days in which *B. breve* NRBB57 displayed metabolic and morphological adaptations in comparison to the chemostat cultivated cells (μ=0.025 h^−1^).

### Viable-but-non-culturable cells

During the 3 weeks of retentostat cultivation, the growth rate of *B. breve* NRBB57 progressively decreased from 0.025 to 0.00092 h^−1^, the latter corresponding to a doubling time of 31 days. Throughout the fermentation, the biomass concentration increased approximately 6-fold. At the same time, culturability significantly reduced to approximately 30%, while viability remained higher than 80% indicating that 20% of the cells were dead while 50% entered a viable but non culturable state. Morphological changes were also progressively observed during the retentostat cultivation showing cell elongation and more branching. These results correspond to what has been observed in *Lactococcus lactis* and *Pseudomonas putida* (10, 11, 18).

In the retentostat, nutritional stress is gradually induced because biomass accumulates while the influx of nutrients remains the same. This nutritional stress can strongly impact not only the morphology of bacteria but also on its culturability (30). The loss of culturability or the increase in the number of cells in the VBNC state in retentostat cultures of *P. putida* has previously been linked to a decrease in ribosome content (18). The latter phenomenon could also be deduced from the proteome data of *B. breve* NRBB57, which showed a lower abundance of ribosomal proteins during the retentostat cultivation indicating a decrease in ribosomal synthesis. Changes in cell morphology have also been linked to VBNC cells as an adaptation to stressful environments. It has previously been hypothesised that this is a mechanism of adaptation to reduce energy requirements (31). By reduced cell division, cells tend to become extended instead of forming a septum and divide (10). This agrees with (i) our observations of a 3.6-fold reduction of the maintenance requirements of *B. breve* after 3 weeks of retentostat cultivation and (ii) morphological changes, as is seen with *L. lactis* as well (Ercan *et al*., 2013; Vos *et al*., 2016; Van Mastrigt *et al*., 2018; Van Mastrigt *et al*., 2019b).

### Stringent response

Proteins involved in the production of the alarmones guanosine tetraphosphate (ppGpp), guanosine pentaphosphate (pppGpp) and guanosine 5′-monophosphate 3′-diphosphate (pGpp) (collectively referred as ppGpp) had higher abundances at the extremely low growth rates which indicates induction of the stringent response. The stringent response is a stress signalling system mediated by (pp)pGpp in response to nutrient deprivation that controls many important processes, such as DNA replication, transcription, ribosome synthesis and maturation (33). The production of (pp)pGpp also plays a key role in inducing the non-growing state in the bacterial cells (34, 35), which could explain why *B. breve* NRBB57 progressively entered a viable-but-non-culturable state during the retentostat cultivation.

The alarmone (pp)pGpp inhibits ribosomal synthesis and maturation, while it up-regulates genes involved in nutrient acquisition (e.g. nutrient transporters) and stress resistance (36). A possible indication of the activation of the stringent response by (pp)pGpp accumulation in our study was observed when 15 out of 52 ribosomal proteins were significantly less abundant at near-zero growth rates compared to at a growth rate of 0.025 h^−1^. In contrast, many amino acid biosynthesis proteins were more abundant at near-zero growth rates as well as several proteins involved in protein and DNA repair. Moreover, 10 different tRNA-synthetases, which load amino acids to tRNAs, were significantly more abundant at near-zero growth rates.

Moreover, proteomics results also showed the up-regulation of polyphosphate kinase at lower growth rates, this enzyme is involved in the production and accumulation of inorganic polyphosphates. Thereby, the stringent response inhibits exopolyphosphatase and triggers the activity of the polyphosphate kinase and subsequent accumulation of polyphosphate (37, 38). Polyphosphate accumulates in response to nutrient starvation as seen in the retentostat cells, and plays an important role in the stress response interacting in several important cellular processes that include inhibiting reinitiation of DNA replication and promoting fitness during starvation (39, 40).

### Pyruvate dissipation

The above observations indicate that in the retentostat *B. breve* NRBB57 down-regulates growth-related processes as a consequence of the reduction in the concentration of lactose available but started to utilize alternative energy sources, like amino acids and glycerol. Moreover, in all retentostat and chemostat cultures pyruvate was mainly converted to acetate, ethanol and formate for extra energy generation. Even at the highest growth of 0.4 h^−1^ with a specific substrate consumption rate of 10 mmol hexose equivalents.gDW^−1^.h^−1^, lactate production was low in the chemostat cultures, while lactose is almost exclusively converted to lactate and acetate in batch cultures of *B. breve* NRBB57 (data not shown). This indicates that the metabolic shift is not only controlled by the specific rate of sugar consumption (41), but also the sugar concentration might play an important role. While lactose was still mainly converted into acetate, formate and ethanol at high growth rates, in the retentostat lactate production increased. Nonetheless, lactate was still the product with the lowest concentration detected in the retentostat. The slightly higher production of lactate in the retentostat could be associated to the significant higher abundance of two L-lactate dehydrogenases observed in our proteomic results.

### Alternative energy sources

In addition to downregulation of growth-related processes to save energy, consumption of alternative nutrient sources could provide the bacteria with valuable energy during the extreme energy limitation imposed by retentostat cultivation. In this study, the concentration of several amino acids gradually decreased during the retentostat cultivation, while ammonia increased accordingly, pointing to a possible role for amino acids as alternative carbon and energy sources. These amino acids include glutamine/arginine, aspartic acid and serine. Because glutamine/arginine depletion coincided with an increase in ornithine, increased arginine utilization was more likely than glutamine utilization.

Although arginine utilisation coincided with ornithine production, no arginine deiminase and/or arginase have been identified in the genome of *B. breve*. Therefore we hypothesize that arginine is catabolised by reversing the arginine biosynthetic pathway (Fig. 9). All three enzymes argininosuccinate lyase (ASL), argininosuccinate synthase (ASS) and ornithine carbamoyltransferase (OTC) have been described to catalyse reversible reactions (42–44). Proteome analysis indicated that both ASS and OTC were significantly more abundant at near-zero growth rates when arginine was catabolised and also ASL and aspartate ammonia lyase (AAL) were present at near-zero growth rates. In many bacteria, the produced carbamoyl-phosphate is further degraded by carbamate kinase (CK) to CO_2_ and NH_3_ yielding ATP, but no gene encoding CK has been identified in *B. breve*, neither in any other *Bifidobacterium* species. Interestingly, both the small and large subunit of carbamoyl-phosphate synthase (CPS) were significantly more abundant at near-zero growth rate. Therefore, it is tempting to speculate that either the CPS catalyses a reversible reaction in *B. breve*, or there is a carbamate kinase in *B. breve* that has not been identified yet. Because ASS converts AMP to ATP and another ATP could be produced by CK, in total 3 ATP could be produced per arginine by this hypothetical pathway (Fig. 9), outlining how arginine could serve as alternative energy source.

Aspartic acid and serine concentration decreased during the retentostat cultivation indicating increased utilization at near-zero growth rates. Serine was most likely converted into pyruvate and ammonia by via serine dehydratase (SD) (45, 46), which was significantly more abundant at near-zero growth rates. Subsequently, pyruvate could be converted into acetate by pyruvate formate lyase (PFL), phosphotransacetylase (PTA) and acetate kinase (ACK) to yield 1 ATP per pyruvate (Fig. 9).

Aspartic acid was most likely converted first to fumarate via either aspartate ammonia lyase (AAL) or via aspartate aminotransferase (AAT), malate dehydrogenase (MDH) and fumarase (Fum) (Fig. 11C). AAL was produced and AAT was even significantly more abundant in the retentostat. However, no MDH and Fum encoding genes have been identified in the genome of *B. breve*, although minor MDH activity was found in other *Bifidobacterium* spp. (47). The resulting fumarate is most likely converted to succinate by succinate dehydrogenase (SDH), of which the abundance of both subunits significantly increased at lower growth rates in the chemostat culture and their abundance remained high in the retentostat cultures. This hypothesis is supported by the observed increase in the succinate concentration over time in the retentostat. Flavin adenine dinucleotide (FAD) is reduced to FADH_2_ by conversion of fumarate to succinate while aspartic acid acts as electron acceptor. This allows for the production of 0.5-1 ATP per aspartic acid because relatively more pyruvate can be converted to acetate instead of ethanol. Succinate production from aspartic acid could also potentially explain the observed succinate production in batch cultures of other *Bifidobacterium* species (41) indicating that those species might also metabolize aspartic acid as alternative energy source.

### Glycerol

In addition to the consumption of several amino acids, surprisingly, also glycerol was consumed in the retentostat cultures. Since genes encoding known enzymes involved in glycerol metabolism are absent in the genome of *B. breve* NRBB57, we analysed corresponding proteomes and noted an increased abundance of glycerol-3-phosphate dehydrogenase (G3PDH) at near-zero growth rates in the retentostat. Therefore, we speculate that glycerol was phosphorylated by an unidentified enzyme to glycerol-3-phophate (G3P), which was subsequently oxidised to dihydroxyacetone phosphate (DHAP) entering the glycolytic pathway. Further conversion to ethanol, would result in a redox neutral pathway in which 1 ATP per glycerol could be produced. We did not observe consumption of glycerol by *B. breve* NRBB57 in batch cultures neither was growth observed under such conditions with glycerol as sole substrate for growth, suggesting specific activation of the putative pathway under extreme energy restriction in retentostat cultivations. Additional research is required to elucidate activation and role of glycerol metabolism in *B. breve*.

Finally, we would like to highlight how this study contributes to our understanding of the ecophysiology of *B. breve*. First, we showed the production of acetate, formate, and lactate at near-zero growth rates. These metabolites can act as substrates for cross-feeding of other beneficial bacteria and for the formation of other short chain fatty acids such propionate and butyrate (48). These fatty acids besides helping to maintain a low pH in the gut inhibiting pathogenic bacteria, conceivably contribute to other health benefits to the host (49, 50). Similarly, succinate is also a substrate for cross-feeding of other microorganisms in the gut and an intermediate in microbial propionate synthesis (51). Lastly, ornithine has been suggested to be involved in the support of the homeostasis in the gut mucosa (52). Our findings show how *B. breve* can adapt its metabolism at lower growth rate while simultaneously producing metabolites that can be of great benefit to the human host.

To conclude, in this study we demonstrated that *B. breve* NRBB57 can be cultivated in the retentostat system at near-zero growth rates, while remaining viable and metabolically active. Metabolite and proteome analysis revealed that the extreme energy restriction induced a multifaceted response including stress defence and stringent response, metabolic shifts and activation of alternative energy producing pathways. It is conceivable that microorganisms encounter near-zero growth rates inducing conditions in a range of natural environments, but the mechanisms used to adapt to these conditions are poorly understood. The use of retentostat cultivations provided insights in how *B. breve* adapts to nutrient scarcity and revealed cellular responses uniquely linked to physiology at near-zero growth rates.

## ACKNOWLEDGMENTS

We thank TKI AgriFood and Danone Nutricia Research for the funding of this project (grant number AF16187). In addition, we give special thanks to dr. Joost Gouw and Gido Jehoel (Danone Nutricia Research) for their technical support and assistance in the analysis of the proteomics data and Marcel Giesbers (Wageningen Electron Microscopy Center, Wageningen University and Research) for their assistance with SEM. Finally we thank our colleague Diego Gallegos for his effort and his support at the beginning of this project.

