## Supplemental figures for "Quantitative physiology and proteome adaptations of *Bifidobacterium breve* NRBB57 at near-zero growth rates"

Running title: Physiology of *B. breve* at near-zero growth rates

Keywords: retentostat, proteomics, chemostat, stringent response, metabolism, bifidobacteria.

**SUPLEMENTARY FIGURES AND TABLES**

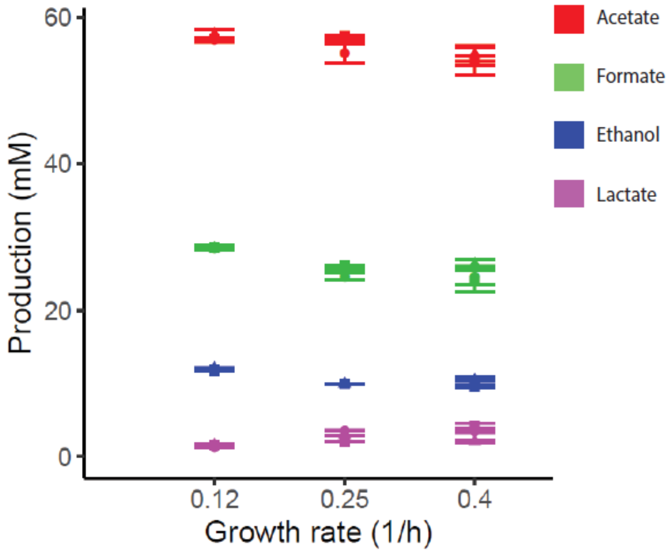

**Figure S1:** Main products of lactose metabolism at diferrent growth rates in the chemostat over time. Colors represent different metabolites, boxplots show the average of three biological replicates. Results showed a constant concentration of acetate, formate, ethanol and lactate.

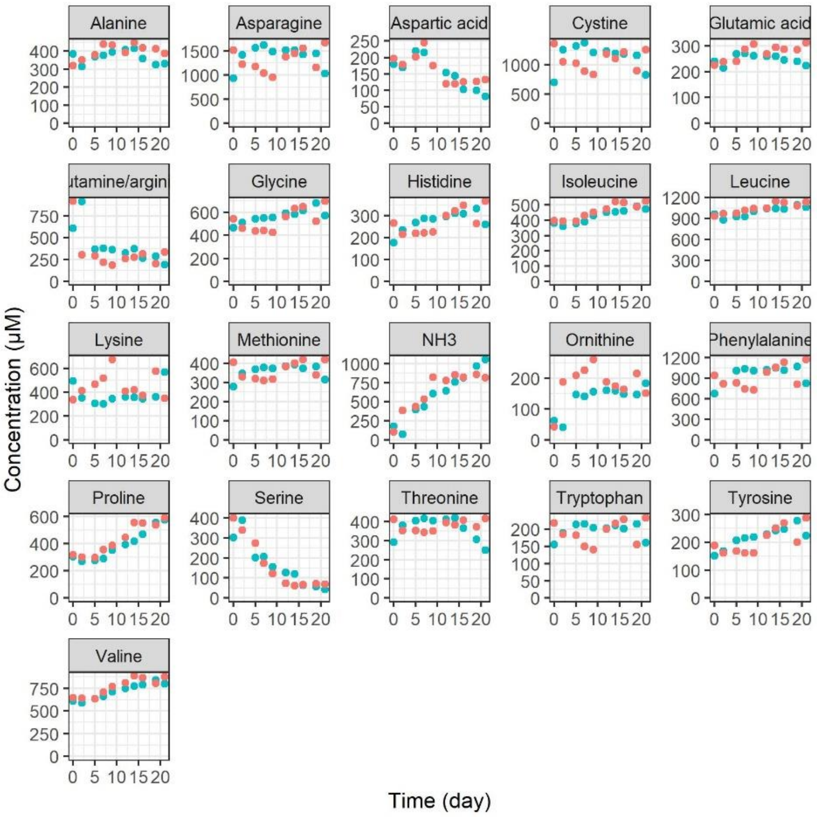


**Figure S2:** Extracellular amino acids and ammonia concentrations in the retentostat measured with UPLC over time. Colors and symbols represent biological triplicates.The concentrations of these amino acids remained stable throughout the retentostat cultivation.

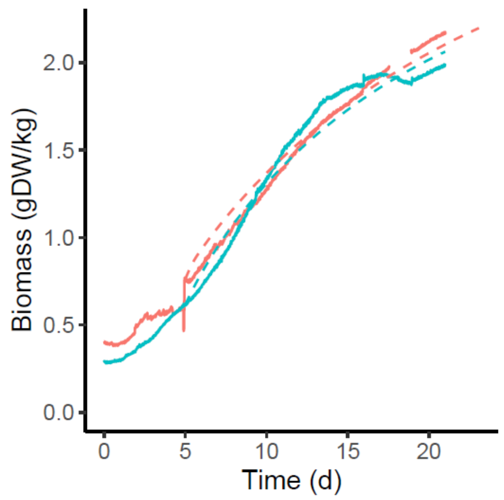


**Figure S3:** Growth of *B. breve* NRBB57 during the retentostat cultivation. Solid lines represent measured biomass accumulation, dashed lines represent the model predictions of the biomass accumulation. The two colors represent 2 different biological replicates.

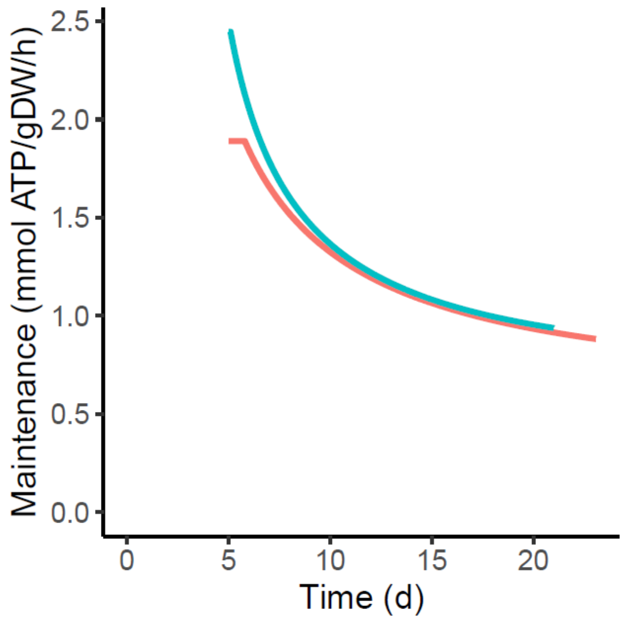


**Figure S4:** Estimated maintenance coefficient of *B. breve* NRBB57 during the retentostat cultivation, starting at day five. The two colour lines represent 2 different biological duplicates. Mathematical explanation has been explained in the experiment procedures.

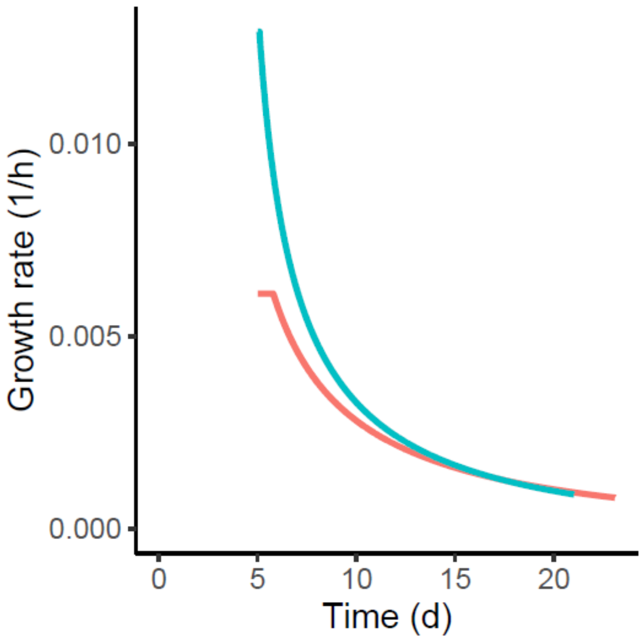


**Figure S5:** Predicted growth rates starting from day five of the retentostat cultivations.

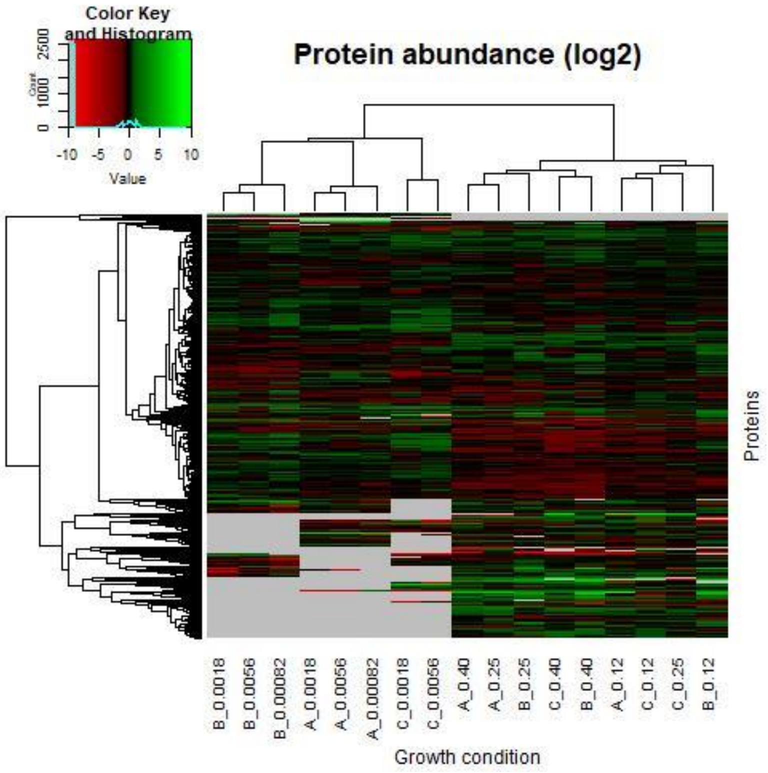


**Figure S6:** Heat map of the proteome of *B. breve* NRBB57 at different growth rates in chemostat and retentostat cultivation. Values are given as log2 ratio compared to the growth rate of 0.025 $h^{-1}$. Green cells represent upregulated proteins, red cells represent downregulated proteins. Grey cells represent proteins that could not be quantified. Labels at the growth condition indicate the replicate (A, B, C) and growth rate.
